# The role of the environment in transmission of antimicrobial resistance between humans and animals: a modelling study

**DOI:** 10.1101/2022.05.19.492687

**Authors:** Hannah C. Lepper, Mark E.J. Woolhouse, Bram A.D. van Bunnik

## Abstract

**Background:** Antimicrobial resistance can be transmitted between animals and humans both directly or indirectly, through transmission via the environment (such as fomites or sewage). However, there is a lack of understanding of, and quantitative evidence about, the contribution of the environment to AMR epidemiology. In this study we incorporate the transmission of resistance via the environment into a mathematical model to study the potential importance of this form of transmission for human resistance levels and any effects of the impact of interventions to reduce antibiotic consumption in animals.

**Methods:** We developed a compartmental model of human-animal AMR transmission with an additional environmental compartment. We compared the outcomes of this model under different human-animal-environment transmission scenarios, conducted a sensitivity analysis, and investigated the impact of curtailing antibiotic usage in animals on resistance levels in humans.

**Results:** Our findings suggest that human resistance levels are most sensitive to both parameters associated with the human compartment (rate of loss of resistance from humans) and parameters associated with the environmental compartment (rate of loss of resistance from the environment and the transmission rate from the environment to humans). The impact of curtailing antibiotic consumption in animals on long term prevalence of AMR in humans was weaker when environmental transmission was assumed to be high.

**Conclusions:** This study highlights that environment-human sharing of resistance can influence the epidemiology of resistant bacterial infections in humans and reduce the impact of interventions that curtail antibiotic consumption in animals. More data on the types and dynamics of resistance in the environment and frequency of human-environment transmission is crucial to understanding the population dynamics of antibiotic resistance.

## Introduction

Antimicrobial resistance (AMR) is a one-health issue, present in a variety of commensal and pathogenic bacteria found in humans, animals and the environment [1],[2]. The potential of the environment for dissemination of AMR has been increasingly recognised, with particular focus on the volume of resistance bacteria in human and agricultural wastewater effluent being discharged into natural waters and soils [3]–[5].

There are many potential routes for resistant bacteria into the environment. Several studies have demonstrated is it likely that resistant bacteria in humans can be transferred to the environment, including rivers[6], coastal waters[7], and soils[8]. In addition, studied have linked resistant bacteria in animals and their respective environments, such as between wild animals and human-impacted environments[9],[10], and between livestock and their environment, especially in aquaculture[11],[12]. However, the risk that the resistance in the environment poses to humans and animals remains poorly understood[13].

Mathematical models are an important tool to study complex dynamics inherent in the emergence and spread of resistance[14] and can therefore be used to improve our understanding and combat the spread of AMR in humans, animals and the environment. However, a lack of data and understanding about the burden, selection and transmission of resistant bacteria, especially in animals and the environment, presents a challenge to parameterising models of AMR from a one-health perspective. Consequently there are few models of resistant bacteria that connect humans, animals and the environment[15].

Some existing studies incorporate an environmental component into transmission models of resistant bacteria in hospitals or farms. Two compartmental models found that reducing or eradicating resistance in a hospital setting was harder to achieve when the environment was explicitly modelled[16],[17]. Studies taking the environment into account when modelling the spread of resistance in farms have found environmental parameters were key in dynamics of resistance in the farm [11],[18]. However, a recent modelling study found that interventions to reduce antibiotic consumption in animals would still be effective when the influence of resistance in animals and the environment is considered[19]. These findings indicate the need for further exploration of the role of the environment with fully dynamic transmission models.

In this study, we aimed to investigate the importance of the environment in the long term dynamics of resistant bacterial infections in humans, including how it might affect the impact of interventions to reduce resistance in humans. A compartmental of resistance transmission within and between humans, animals and the environment was developed. We use a dynamic environmental compartment, improving on existing models by allowing us to assess the importance of within-environment processes. Our objectives were: 1) to investigate how adding an environmental compartment affects the long-term dynamics of resistance in humans, and the sensitivity of the model to its parameters; and 2) to investigate the impact of interventions to curtail antimicrobial usage in animals or environment to human transmission on the prevalence of resistance in humans in this model.

## Methods

### Model description

We extended the original model presented in van Bunnik and Woolhouse, 2017[20], to include an environmental compartment. Humans and animals gain resistant infection by exposure to antibiotics, or exposure to other humans, animals or environments carrying resistant bacteria. Resistance in the environmental compartment is increased by contact with humans or animals who carry resistant bacteria, or via exposure to antibiotics that have been excreted by humans or animals. The environment is not considered to be any one type of environment, such as water or soil, but rather a summation of these types.

We define the model using a system of coupled ordinary differential equations:

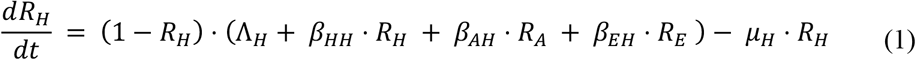

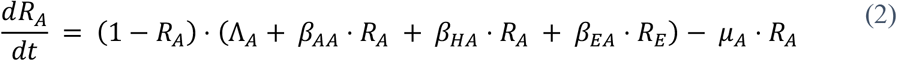

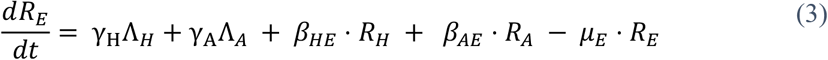

*R*_*H*_ and *R*_*A*_ are the fractions of the human and animal population that are infected with resistant bacteria, respectively, and *R*_*E*_ is a measure of the amount of resistant infectious bacteria in the environment. *Λ*_*H*_ is the constant rate at which resistance is gained from exposure to antibiotics in humans, and Λ_*A*_ is the equivalent in animals. These are composite variables, taking into account both the amount of antibiotics consumed and the rate at which selection causes resistance in bacteria to arise. *µ*_*H*_ is the reversion rate of humans infected with resistant bacteria to having only sensitive bacteria, and *µ*_*A*_ is the equivalent in animals. This includes the rate of clearance of resistant infection and the rate of death in a fixed-size population. The parameters *γ*_*H*_ and *γ*_*A*_ are scaling parameters determining how much of the antibiotic exposure in humans (Λ_*H*_) and animals (Λ_*A*_) will result in excreted antibiotics selecting for an increase in resistant bacteria in the environment. *µE* is the rate of loss of resistant infectious bacteria from the environment. Transmission within and between the compartments is controlled by the *β* transmission coefficients, with the subscripts indicating the direction of transmission of each coefficient. For example, *β*_*HH*_ is the transmission coefficient between humans, and *β*_*EH*_ is the transmission from the environment to humans.

Further details about parameter definitions, units and values ranges can be found in the Appendix Table 1. Fig. 1A shows a flow diagram representing the movement of infectious resistant material between and within the different compartments. All rates are per capita with respect to the human and animal populations, and per environmental unit with respect to the environment (see next section). We used the steady state solutions of this model, obtained numerically, as we were interested in long-term prevalence. The timestep of the model represents one month.

**Figure 1.**
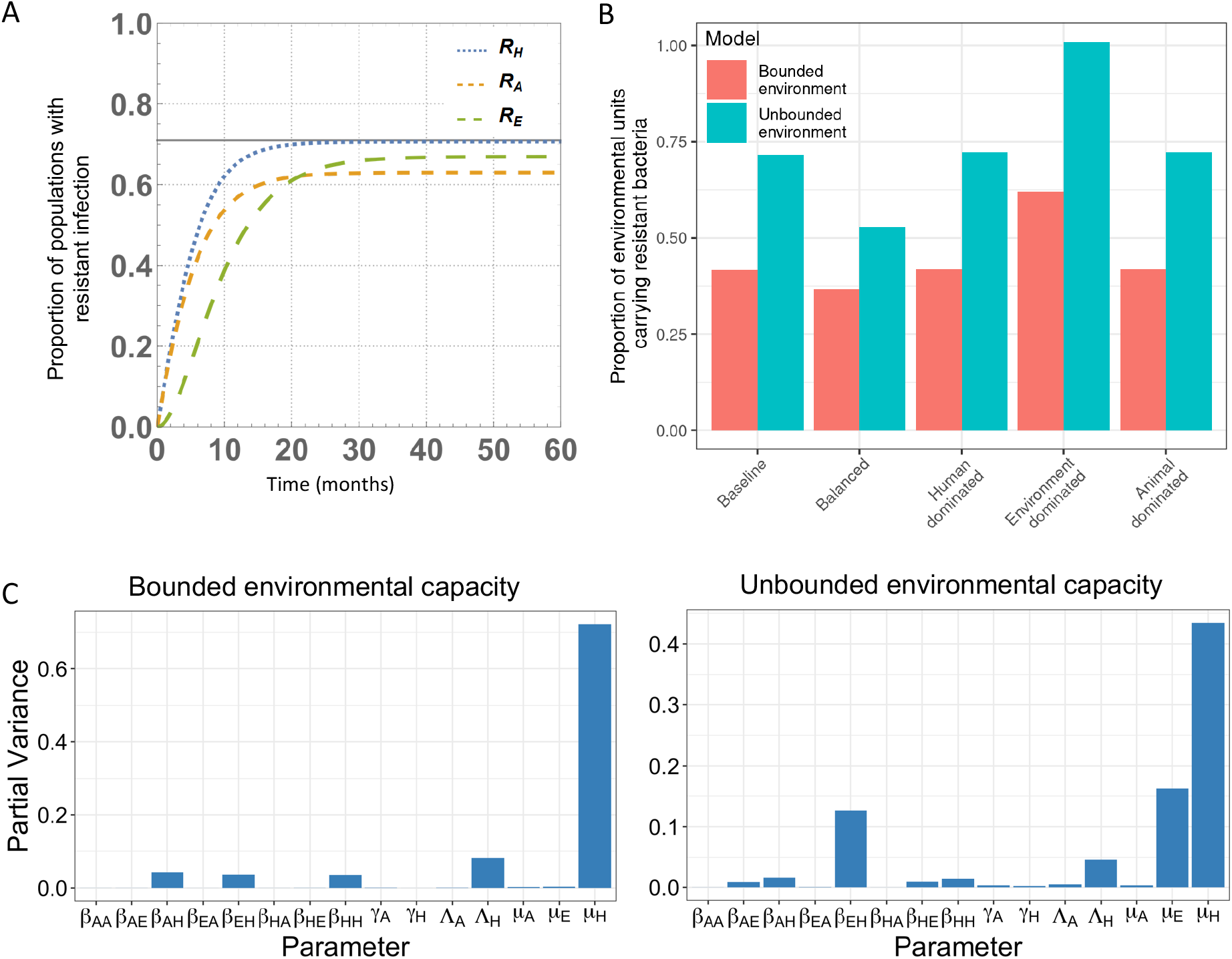
A: flow diagram indicating the model structure. B: *R*_*E*_ values in all transmission scenarios and both model structures. C: Fourier Amplitude Sensitivity Tests (FAST), indicating how much variation in *R*_*H*_ was explained by each model parameter. On the left, FAST for the version of the model in which *R*_*E*_ is bounded to 1. On the right, FAST for the version of the model in which *R*_*E*_ was unbounded.

### Capacity for resistance in the environment

Equation (3) represents the environment as an unbounded compartment, in which the amount of resistant infectious material in the environment is in the range 0 - ∞. We consider one “unit” of the environment to be the human infectious potential equivalent. This means that for a value of *R*_*E*_ = 1, if the transmission coefficients *β*_*EH*_ and *β*_*HH*_ were the same, each unit of the environment would transfer resistant material to humans at the same rate that an infected human would to another human. Although theoretically the environment may have some maximum capacity for resistant material, we do not have a way to determine this capacity, so we modelled the environment as an unbounded compartment. For comparison, we also explored a version of the model in which resistance levels in the environment cannot exceed 1. In this model the environmental compartment is specified:

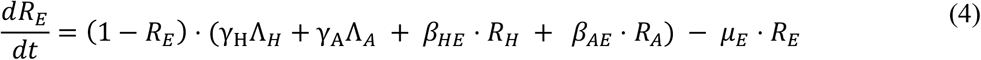

This model assumes that there is no growth or dissemination of resistant organisms within the environment. We also assume that the environment is only exposed to antibiotics that are excreted by humans or animals. The environment may be exposed to antibiotics directly through, for example, the effluent of pharmaceutical industries, but we do not consider this specific case here.

### Impact of interventions on resistance in humans

We investigated the impact of two types of interventions on the levels of resistance in the human compartment. Firstly, we looked at interventions to remove antibiotic usage in livestock (reducing Λ_*A*_ to 0), and how changes to environmental parameters affect the effectiveness of this intervention. Secondly, we looked at interventions that would reduce the transmission of resistant bacteria from the environment to humans (reducing *β*_*EH*_ to 0).

We measured the impact of interventions as the percentage decrease in resistance levels in humans, following van Bunnik and Woolhouse (2017). We compare equilibrium values of *R*_*H*_ before 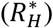 and after the intervention 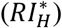, to obtain the impact, or percentage decrease in human resistance levels:

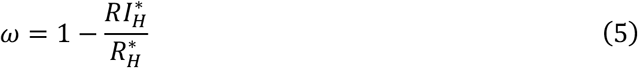

We investigate the impact of reducing *β*_*EH*_ and of curtailing antibiotic usage in animals (Λ_*A*_).

### Sensitivity analysis

We use the extended version of the Fourier Amplitude Sensitivity Test (FAST)[21] to analyse the relative influence of each parameter on the value of *R*_*H*_, the outcome measure of interest. A total sensitivity index for each parameter is calculated based on the variance of *R*_*H*_ over variation in all parameters. The R package fast was used for this analysis[22].

### Parameterisation

Due to a paucity of data about many of the parameters in the model, we aimed to explore a wide range of parameter scenarios in this model. We chose the following transmission scenarios: 1) a baseline, with transmission parameter values similar to those of the original[20]; 2) a balanced transmission scenario, with all transmission coefficients equal; 3) human-driven transmission (i.e., if the subscript H denotes the humans and *x* denotes any other compartment *β*_*Hx*_ > *β*_*xx*_); 4) animal-driven (*β*_*Ax*_ > *β*_*xx*_); and finally 5) environment-driven (*β*_*Ex*_ > *β*_*xx*_).

We also averaged our results across parameter sets generated randomly using sampling distributions for the three parameters *R*_*H*_ that was most sensitive to (viz. *µ*_*H*_, *µ*_*E*_, and Λ_*H*_), to avoid over-reliance on model dynamics that are unusual to a particular combination of parameters rather than generally true of the system. All parameter values and sampling distributions can be found in the Appendix (Tables 2A and 2B), as well as the methods for obtaining transmission scenario parameters.

### Software

Analyses were carried out using Wolfram Mathematica version 11.3[23], R 4.1[24], and Julia 1.7[25]. The code for the model, parameter set generation, and visualisations is available at https://github.com/hannahlepper/animal-human-env-model.

## Results

All analyses were conducted in both bounded and unbounded environmental capacity versions of the model.

### Long term dynamics of resistance in humans

#### Prevalence of resistance in humans

For all transmission scenarios, parameter sets were identified that corresponded to the intended target equilibrium human resistance prevalence of 71% in both the bounded and unbounded versions of the model (Appendix Fig. 1). Fig. 1B shows that the amount of resistance in the environment was influenced by the model structure and the transmission scenario. The highest level of resistance in the environment was in the environment-driven, unbounded version of the model, indicating that an implausibly high level of environmental contamination is not needed for observed human resistance levels.

#### Sensitivity analysis

Model sensitivity results are presented in Fig. 1C. In both bounded and unbounded models, human resistance prevalence was most sensitive to *µ*_*H*_, the rate of loss of resistance from humans, but relatively insensitive to *Λ*_*A*_, the antibiotic consumption in animals. The rate of transmission from the environment to humans, *β*_*EH*_, was at least as important as *β*_*HH*_ and *β*_*AH*_, rates of transmission to humans from other humans and from animals. Moreover, *β*_*EH*_ is more influential than any other transmission parameter in the unbounded model. The rate of loss of resistance from the environment, *µ*_*E*_, was more important for human resistance levels in the unbounded than the bounded model.

### Impact of interventions to reduce resistance in humans

#### Impact of curtailing antibiotic usage in animals

Curtailing antibiotic usage in animals had a small impact on human resistance levels, and the impact was lower when the environment was explicitly modelled or when animals contributed less to resistance transmission (Fig 2). The percentage decrease in human resistance levels achieved *without* an environmental compartment and using the parameters of the original model (the ‘baseline transmission scenario) was 3.2%. Simply adding an environmental compartment and keeping other parameters reduced the percentage decrease to 2.8% in the unbounded and 2.9% in the bounded model. The animal-driven transmission scenario had the highest impacts (5.8% decrease in human prevalence), and the human-driven scenario had the lowest (0.064%). In the environment-driven transmission scenario, the environmental capacity was influential: when bounded, the impact was low (0.94%), and increased when unbound (3.2%). Both the environmental structure and the transmission parameters affected the impact of antibiotic usage reduction in animals.

**Fig 2:**
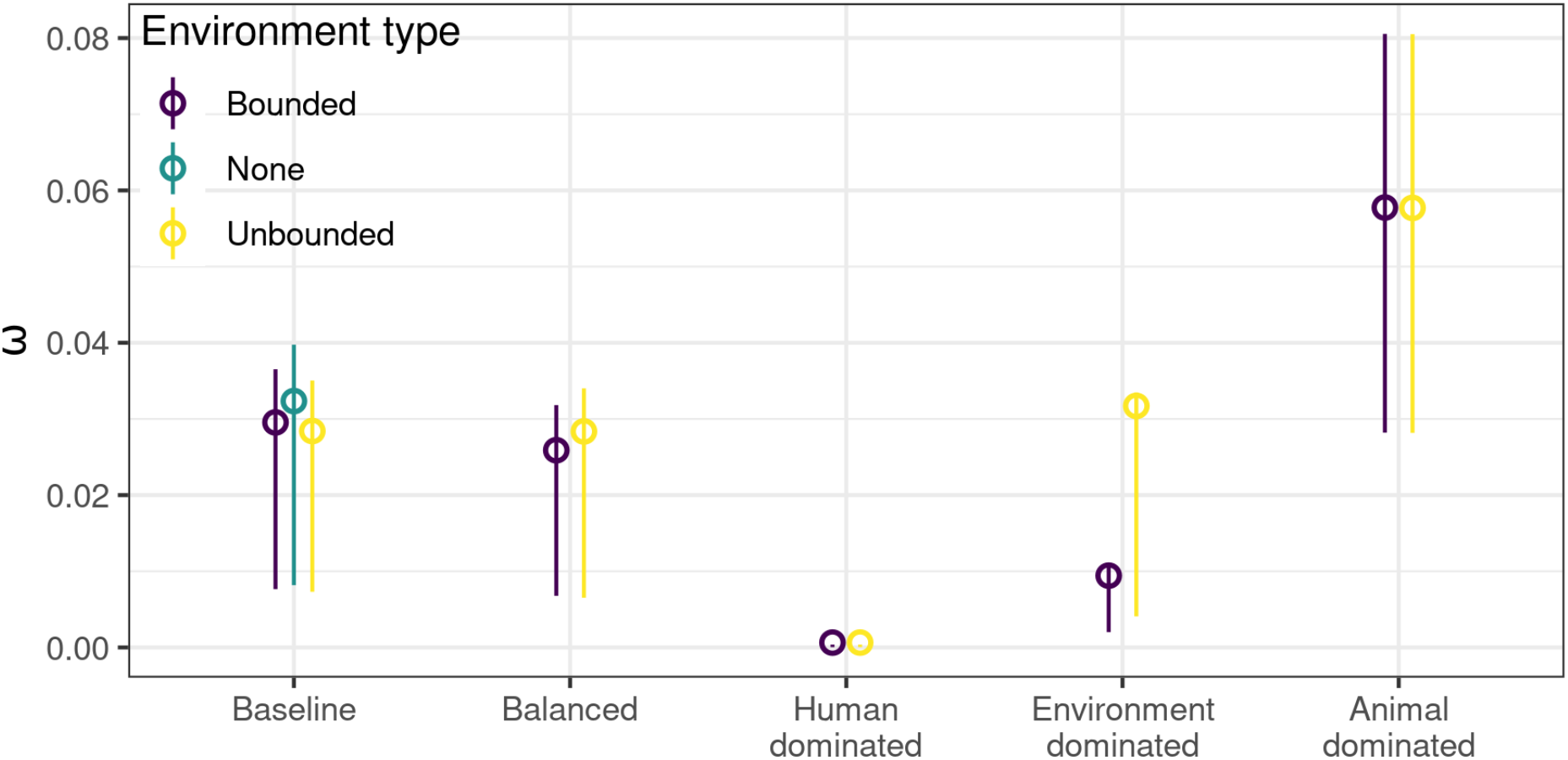
Mean impact of reducing Λ_*A*_ from 0.1 to 0 across transmission scenarios. The green point in the baseline transmission scenario group is the mean impact for the original van Bunnik and Woolhouse (2017) model, with no environmental compartment included. Results were averaged for parameter sets with *µ*_*H*_, *µ*_*E*_, and Λ_*H*_ varied, with error bars indicating standard deviation in results.

#### *Reducing Λ*_*A*_ *vs. reducing β*_*EH*_

We compared the impact (ω) of reducing either Λ_*A*_ (the antibiotic consumption in animals) or *β*_*EH*_ (the transmission of resistant material from the environment to humans) (Fig. 3). We considered pre-intervention values of 0.1 for each parameter, as well as the impacts in different transmission scenarios. This value was chosen so that the size of the intervention was consistent between transmission scenarios in this model and with the previous model (van Bunnik and Woolhouse, 2017). Interventions to reduce *β*_*EH*_ had a greater impact than interventions to curtail Λ_*A*_ when transmission was human- or environment-driven, or when transmission was balanced. When livestock dominated transmission or for the baseline parameter set, the impacts of interventions to reduce *β*_*EH*_ or Λ_*A*_ were similar.

**Fig 3.**
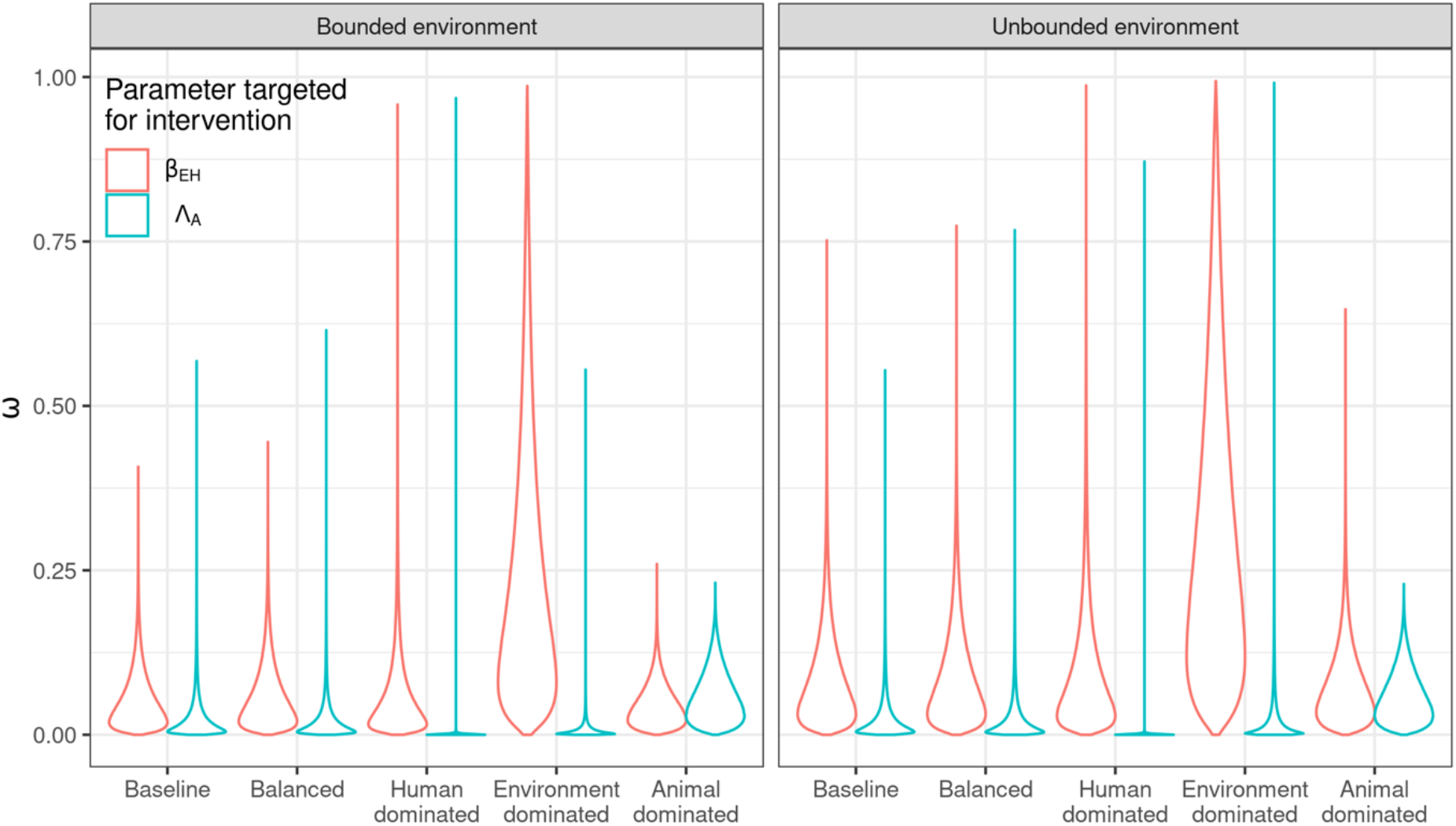
Violin plots of the impact (proportion decrease in *R*_*H*_ after the intervention) of reducing either *β*_*EH*_ or Λ_*A*_ in all transmission scenarios and for both model structures. The intervention target was reduced from 0.1 to 0 in each case for consistency.

#### Effect of β_EH_ on impact of interventions to reduce antibiotic consumption in animals

We next identified the impact of reducing Λ_*A*_ across a range of values for *β*_*EH*_ (Fig. 4). Increasing *β*_*EH*_ decreased the size of the impact of curtailing antibiotic usage in animals in all transmission scenarios (Fig 4A). The peaked shape of the impact size in the environmental transmission scenario is caused by the increase in *β*_*EH*_ allowing increasing indirect transmission in animals and humans. This effect is only observed when there is little non-environmental transmission. Fig. 4B shows that the decrease in intervention impact was also observed across the range of pre-intervention values for Λ_*A*_. These results indicate that increasing environmental transmission can reduce the impact of curtailing antibiotic usage in animals.

**Fig 4:**
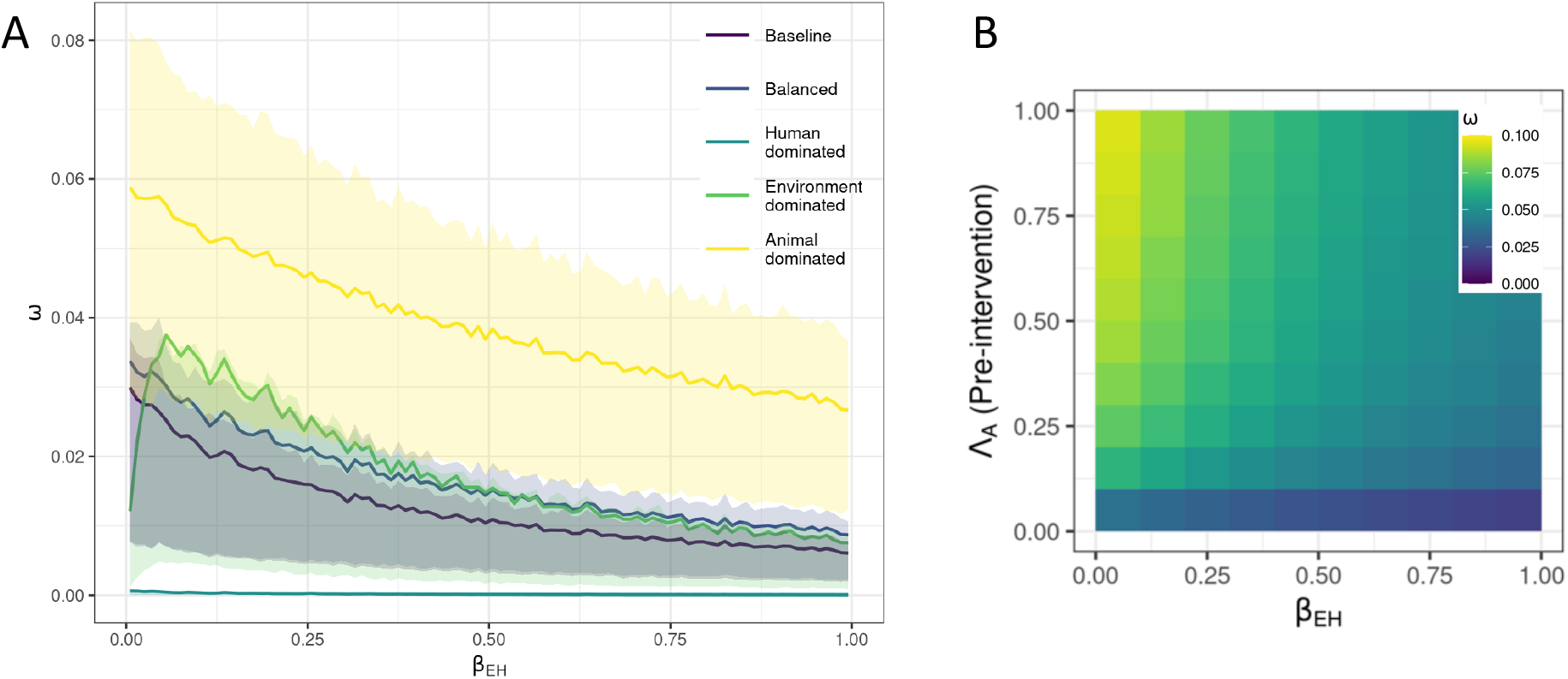
A) Mean impact of antibiotic decrease in animals on human resistance levels (proportion decrease in human resistance levels) for each transmission scenario with increasing rate of environment to human transmission (*β*_*EH*_). Ribbons indicate 25% and 75% impact quantiles. B) Heatmap of the impact of different pre-intervention values of Λ_*A*_ (y axis) against different levels of environment to human transmission, *β*_*EH*_ (x axis), for the animal transmission scenario in the unbounded model. The colour of the tiles indicates the average value of the impact of the intervention from 17,000 parameter sets where *µ*_*H*_, *µ*_*E*_, and Λ_*H*_ were varied.

## Discussion

### Key findings

In this study we modelled the transmission of resistant bacteria between humans, livestock animals and the environment, and assessed the impact of interventions that reduce antibiotic consumption in animals or decrease the transmission of resistant bacteria from the environment to humans. We found that human resistance prevalence is sensitive to transmission between humans and the environment. Including an environmental compartment in the model decreased the impact of curtailing antibiotic resistance, and a more transmissible environmental reservoir of resistant bacteria further mitigated the impact of this intervention. Reducing the transmission of resistant bacteria from the environment to humans was found to be a more effective intervention than reducing antibiotic consumption in animals. Overall, these results indicate that resistant bacteria in the environment can influence the prevalence of resistance in humans. The size of environmental influence will depend on the amount and dynamics of resistant bacteria in the environment. Assessing the likelihood of observing these theoretical results in the real world is hindered by a lack of quantified, generalisable data on the types, amount, and degradation of resistance in the environment, and the transmission of resistance between humans, livestock and the environment.

### Is curtailing antibiotic usage in animals an effective intervention to reduce human resistance levels?

The greatest observed impact of curtailing antibiotics in animals was a modest 10% decrease in human resistance level in a balanced transmission scenario, and the smallest impact was a <1% in the human-driven transmission scenario. This result provides little theoretical support that curtailment of antibiotics would appreciably decrease resistance in humans in many settings. In contrast, there is some empirical evidence that curtailing antibiotics in livestock could reduce human resistance levels, although from a small set of observational studies[26]. A study of use of third-generation cephalosporin ceftiofur in broiler rearing in Canada found that resistance in *Salmonella* and *E. coli* was decreased in clinical isolates by 20% and 40%, respectively, after ceftiofur use decreased[27]. This real-world population-level effect is greater than our results would predict, and may indicate they are an underestimate, especially with respect to the degree of sharing of resistance between humans and animals. More data-based parameterisation will be crucial to improve the accuracy of one-health resistance transmission models.

The size of the effect of intervening to reduce antibiotic consumption in livestock varied by transmission scenario (balanced transmission, or transmission driven by either humans, livestock or the environment). Therefore, a key question for assessing the accuracy and relevance of the resulting intervention effect sizes is how realistic are the transmission scenarios? Although transmission of resistance between humans and animals is of great concern, evidence that conclusively demonstrates a case of direct transmission is rare[28],[29]. Accurately parameterising the relationship between resistance in humans and livestock is an ongoing area of research[30] which will be crucial for one-health modelling of resistance. It seems likely that on average across a large human population, human-human transmission is far more common than animal-human transmission and we suggest human-driven scenario to be most relevant for resistance dynamics in the human population.

As we increased the transmission rate from the environment to humans, the effectiveness of antibiotic curtailment was decreased. This suggests that the environment can provide a ‘back door’ transmission route from animals to humans that can reduce the effectiveness of antibiotic curtailment by adding to overall animal-human transmission rates. Using a two-pronged approach by intervening to reduce environmental transmission at the same time could therefore improve the impact of antibiotic usage curtailment. However, the effect of environmental transmission on antibiotic curtailment effectiveness was negligible in the human-dominated transmission scenario (Appendix Fig. 2.), again indicating the importance of transmission setting for this result. It remains unclear if non-human dominated transmission scenarios are realistic, and therefore what the real-world size of this back-door effect might be. There is some evidence that microbiomes in humans, animals and the environment become more shared as interactions become more frequent[31], suggesting that transmission scenarios in which humans do not dominate transmission (such as the balances and baseline scenarios) are possible. Further work to quantify environmental resistance concentrations and transmission could improve accuracy of outcome predictions of antibiotic usage interventions. As reducing antibiotic usage in livestock animals is a costly intervention, it is important to ensure optimal implementation.

### Could the environment be an effective alternative intervention target?

The rate of transfer of resistant bacteria from environment to humans (*β*_*EH*_) is also a potentially effective intervention target. Human resistance prevalence levels were sensitive to *β*_*EH*_ and *µ*_*E*_, the rate of loss of resistant bacteria from the environment (sensitivity analysis, Fig. 1C), which suggests that interventions to reduce how much resistance humans gain from the environment would be effective. Indeed, the impact of reducing *β*_*EH*_ was more effective than antibiotic usage curtailment interventions, although the difference was small in the animal-dominated scenario (Fig. 2A). Interventions that improve sanitation have been proposed to reduce occurrences of transmission of resistance between humans and the environment in informal urban communities in LMICs where there is frequent exposure to resistance bacteria in the environment[32],[33]. Nadimpalli et al (2020) focus particularly on the potential benefits of improved water and wastewater infrastructure for controlling and preventing AMR transmission, but note that few studies have investigated the impacts of sanitation interventions on AMR.

### Should the environment be included in AMR models?

In this model, the environment played an important role in the long-term dynamics of antibiotic resistance levels in humans. Mechanistically, the environment acts as a reservoir for antibiotic resistance from humans and animals in this model structure. Therefore, parameters that provide more opportunity for transmission to humans were influential in human resistance levels, especially the rate of loss or level of persistence of resistant bacteria in the environment (*µ*_*E*_). Environmental parameters were also influential in the size of impact of interventions, and we show that it may be an effective intervention target itself. Existing models that incorporate an environmental component have also highlighted the potentially strong role the environment could play in increasing resistance levels in humans and undermining interventions[16]–[19]. Most models include environment as a constant rather than a dynamic compartment, with the exception of Booton et al, 2021. As we find comparable results to models with constant compartments, this may indicate that models incorporating the environment simply may be enough to account for this additional source of resistant bacteria. On the other hand, the model in Booton et al, 2021, assumes that transmission of resistance (including from the environment) is dependent on exposure to antibiotics and accordingly finds that human antibiotic usage is the most influential parameter for human resistance, downplaying the role of the environment. This contrasting result points to a need for further models that compare the contribution of the environment under different model structures and assumptions. Incorporating the environment into models of AMR spread may be important in understanding AMR prevalence and for evaluating intervention success.

### Modelling the environment highlights data needs

The results highlight some key data needs for understanding the importance of AMR in the environment for humans. There are two influential parameters in the model which are difficult to parameterise from existing data: the rate of transfer of AMR from the environment to humans, and the rate of loss of resistance in the human population.

How frequently humans gain resistant bacteria after exposure to an environmental source is unknown. There is evidence that humans can be exposed to resistant bacteria in the environment. For example, one study estimated that the amount of third-generation cephalosporin resistant *E. coli* that humans would ingest during recreational water use in coastal regions in England and Wales poses a risk of infection[7]. However it is not clear how often these exposures lead to infection or colonisation[34]. More research that demonstrates a close relationship and epidemiological link between resistant bacteria colonising the environment and humans is needed to understand the frequency of environment-human transmission events. Use of high resolution typing such as whole genome sequencing of, for example, isolates from hospital patients and the hospital environment in longitudinal studies would be ideal for this research.

Studies have provided data on the rate of clearance of resistant infections in humans. A systematic review on methicillin-resistant *S. aureus* (MRSA) and vancomycin-resistant *Enterococcus* (VRE) colonisation found that it takes a period of 88 and 26 weeks on average to clear MRSA and VRE infections, respectively[35]. However, they note that there is considerable methodological heterogeneity in studies of MRSA and VRE, including varying definitions of clearance, and length of follow-up[35]. The studies also focussed primarily on hospital-associated resistance. Data on resistant bacteria colonisation prevalence and clearance in the community, where the role of exposure to animals and the environment may play a greater role, appear to be rare. Parameterising generalisable one-health models will therefore be benefitted by more research into resistance in the community.

### Limitations

There are some important limitations to this study that should be noted. Firstly, we make simplifying assumptions in the structure and parameterisation of the model. These are suitable to the questions posed in this study, but there are still many complexities in the spread and emergence of AMR in humans, animals and the environment to be explored. Further models should explore the importance of potential complexities, such as heterogeneity of transmission events, separate humans-specific and animal-specific environmental reservoirs, variation in the capacity for resistance in the environment, or the fitness costs to bacteria of carrying resistance in the three populations.

We do not model the dynamics of transmission of resistant bacteria and resistance genes separately, but assume that transmission parameters combine the transmission of both. This is in-keeping with the assumptions of the original model [20]. Resistance genes can be transferred between bacteria via plasmid transfer or bacteriophages, and can also be lost from bacterial lineages. The transmission rates of resistance genes in human population may therefore differ from resistant bacteria, and it is a limitation that we do not capture this in the model. AMR epidemiology and surveillance is usually measured in resistant bacteria so there is little data on the prevalence and transmission rates of specific resistance genes.

Two further assumptions about resistance in the environment are that we assume that there is no growth of resistant material within the environment, and that all antibiotics secreted into the environment are from human and livestock usage. The dynamics of resistance genes and bacteria in the environment is a complex topic, and although there are potentially environments in which resistance may spread (especially in sewage) much more empirical and modelling research is needed[34],[36]. A recent review found that the sources of antibiotics in ground water include excretion from humans and animals (via sewage and manure) but also landfill, aquaculture and industrial sites[37], so not including these sources may limit the accuracy of the results of this model. However the relative contribution of each sources is not well known and may vary from one country to another[37].

## Conclusions

This study illustrates the potentially important role of the environment in the epidemiology of resistant bacterial infections in humans. We highlight the need to consider the role of the environment in the design of AMR control strategies, as it can be influential in human prevalence of resistance, reduce the effectiveness of interventions that curtail antibiotic consumption in animals, and may be an effective intervention target itself via improved sanitation infrastructure. Incorporating the environment into a one-health model of antibiotic resistance as a dynamic compartment was useful for considering the role of the environment. However, assessing the uncertainty of model predictions is hindered by a lack of data on the types and frequency of resistance in the environment, and the frequency of environment-human transmission events.

