## Supplementary Materials for "The role of the environment in transmission of antimicrobial resistance between humans and animals: a modelling study"

#### Table 1: Parameter definitions

| **Parameter** | **Definition and units** |
| --- | --- |
| $\beta_{HH}$ | Per capita rate at which humans acquire antibiotic resistant bacteria as a result of exposure to other humans harbouring resistant bacteria per time step |
| $\beta_{AA}$ | Per capita rate at which animals acquire antibiotic resistant bacteria as a result of exposure to other animals harbouring resistant bacteria per time step |
| $\beta_{AH}$ | Per capita rate at which humans acquire antibiotic resistant bacteria as a result of exposure to animals harbouring resistant bacteria per time step |
| $\beta_{HA}$ | Per capita rate at which animals acquire antibiotic resistant bacteria as a result of exposure to humans harbouring resistant bacteria per time step |
| $\beta_{HE}$ | Per environmental unit rate* at which the environment acquires resistant bacteria as a result of exposure to humans harbouring resistant bacteria per time step |
| $\beta_{EH}$ | Per capita rate at which humans acquire antibiotic resistant bacteria as a result of exposure to environmental units harbouring resistant bacteria per time step |
| $\beta_{AE}$ | Per environmental unit rate* at which the environment acquires resistant bacteria as a result of exposure to animals harbouring resistant bacteria per time step |
| $\beta_{EA}$ | Per capita rate at which animals acquire antibiotic resistant bacteria as a result of exposure to environmental units harbouring resistant bacteria per time step |
| $\Lambda_{H}$ | Per capita rate at which humans acquire antibiotic resistant bacteria as a result of direct exposure to antibiotics per time step |
| $\Lambda_{A}$ | Per capita rate at which animals acquire antibiotic resistant bacteria as a result of direct exposure to antibiotics per time step |
| $\gamma_{H}$ | Proportion of $\Lambda_{H}$ that reaches the environment as antibiotics (a scalar parameter) |
| $\gamma_{A}$ | Proportion of $\Lambda_{A}$ that reaches the environment as antibiotics (a scalar parameter) |
| ${\gamma_{H}\Lambda}_{H}$ | Per environmental unit rate* at which the environment acquires resistant bacteria as a result of exposure to a proportion of antibiotics given to humans per time step |
| $\mu_{H}$ | Per capita rate at which humans with resistant bacteria revert to have only sensitive bacteria per time step |
| $\mu_{A}$ | Per capita rate at which animals with resistant bacteria revert to have only sensitive bacteria per time step |
| $\mu_{E}$ | Per environmental unit rate* at which environmental units harbouring resistant bacteria revert to having only sensitive bacteria per time step |

#### Table 2: Parameter values

##### Table 2A: Transmission coefficients

Unbounded model

| **Parameter** | **Value** | | | | | |
| --- | --- | --- | --- | --- | --- | --- |
|  | Baseline | Balanced | Balanced (low $\beta_{HA}$) | Environment-driven | Animal-driven | Human-driven |
| $\beta_{HH}$ | 0.1 | 0.07432092 | 0.07432092 | 0.001 | 0.001 | 0.2019663 |
| $\beta_{AA}$ | 0.1 | 0.07432092 | 0.07432092 | 0.001 | 0.2019663 | 0.001 |
| $\beta_{HA}$ | 0.1 | 0.07432092 | 0.00074321 | 0.001 | 0.001 | 0.2019663 |
| $\beta_{AH}$ | 0.1 | 0.07432092 | 0.07432092 | 0.001 | 0.2019663 | 0.001 |
| $\beta_{EH}$ | 0.01 | 0.07432092 | 0.07432092 | 0.1420501 | 0.001 | 0.001 |
| $\beta_{EA}$ | 0.01 | 0.07432092 | 0.07432092 | 0.1420501 | 0.001 | 0.001 |
| $\beta_{HE}$ | 0.1 | 0.07432092 | 0.07432092 | 0.1420501 | 0.001 | 0.2019663 |
| $\beta_{AE}$ | 0.1 | 0.07432092 | 0.07432092 | 0.1420501 | 0.2019663 | 0.001 |

Bounded model

| **Parameter** | **Value** | | | | | |
| --- | --- | --- | --- | --- | --- | --- |
|  | Baseline | Balanced | Balanced (low $\beta_{HA}$) | Environment-driven | Animal-driven | Human-driven |
| $\beta_{HH}$ | 0.1 | 0.08109928 | 0.08109928 | 0.001 | 0.001 | 0.20239149 |
| $\beta_{AA}$ | 0.1 | 0.08109928 | 0.08109928 | 0.001 | 0.20239149 | 0.001 |
| $\beta_{HA}$ | 0.001 | 0.08109928 | 0.00081099 | 0.001 | 0.001 | 0.20239149 |
| $\beta_{AH}$ | 0.1 | 0.08109928 | 0.08109928 | 0.001 | 0.20239149 | 0.001 |
| $\beta_{EH}$ | 0.01 | 0.08109928 | 0.08109928 | 0.23084954 | 0.001 | 0.001 |
| $\beta_{EA}$ | 0.01 | 0.08109928 | 0.08109928 | 0.23084954 | 0.001 | 0.001 |
| $\beta_{HE}$ | 0.1 | 0.08109928 | 0.08109928 | 0.23084954 | 0.001 | 0.20239149 |
| $\beta_{AE}$ | 0.1 | 0.08109928 | 0.08109928 | 0.23084954 | 0.20239149 | 0.001 |

##### Table 2B: Other parameters

| **Parameter** | **Value** | |
| --- | --- | --- |
|  | **Fig 1. B** | **Fig 2. And 3.** |
| $\Lambda_{H}$ | 0.1 | Beta(1.7, 15.3) (mean 0.1) |
| $\Lambda_{A}$ | 0.1 | No intervention: 0.1 or U(0.000001,1.);  intervention: 0.0. |
| $\gamma_{H}$ | 0.001 | 0.001 |
| $\gamma_{A}$ | 0.001 | 0.001 |
| $\mu_{H}$ | 0.1 | Beta(1.7, 15.3) (mean 0.1) |
| $\mu_{A}$ | 0.1 | 0.1 |
| $\mu_{E}$ | 0.2 | Beta(3, 12) (mean 0.2) |

### Methods for finding transmission parameter coefficients

Transmission parameters were chosen by the following method: some parameters were fixed ($p_{f}$) whilst the transmission parameters of interest varied ($p_{v}$) to reach a human resistance level of 71% (prevalence of ampicillin resistance in the UK in 2010, ECDC, 2010):

$$\min_{p_{v}\in\left( 0,1 \right)} \left| 0.71-f\left( p_{v}, p_{f} \right) \right|$$

For example, for the human-driven transmission scenario, rates of transfer of resistance from humans to any other population were varied ($\beta_{HH}=$ $\beta_{HA}=$ $\beta_{HE}$), and all other transmission parameter values were fixed at a low value, 0.001.

### Model timesteps

We selected a value of 0.2 for $\mu_{E}$ in the baseline set of parameters. Zhang et al, 2017 estimated for the rate of loss of various resistance genes from a compost microcosm experiment was 0.0077 per day. We can therefore estimate that the units of our select value is per ~26 days (0.2/0.0077). We intend this model to be used for looking at long term prevalence of resistance in humans, so this estimate of time step is reasonable as it allows us to look over the timescale of years.

To ensure equilibrium values were obtained for all experiments, we initially numerically solved the model to 500 timesteps, and if there was a difference of more than 0.0000001 between the $R_{H}$ values for the final two timesteps, we solved to 10,000 timesteps.

### Bounded model

To investigate the impact of the assumption that the environment has no carrying capacity for resistant organisms, we compared the results of the unbounded model with a bounded version. In the bounded version of the model, we replace the equation for $\frac{dR_{E}}{dt}$ with:

$$\begin{aligned} \frac{dR_{E}}{dt}=\left( 1-R_{E} \right)\left( {\gamma{}_{H}\Lambda}_{H}+{\gamma{}_{A}\Lambda}_{A} + \beta_{HE}\cdot R_{H} + \beta_{AE}\cdot R_{A} \right) - \mu_{E}\cdot R_{E} \# \end{aligned}$$

Of the results comparisons made, the greatest difference was in the sensitivity analysis, which can be found in Fig. 1. C. in the main text.

Figure 1: Trajectory plot of the fraction of human and animal populations carrying resistant bacteria ($R_{H}$, $R_{E}$), and the amount of resistant material in the environment ($R_{E}$). For bounded environment model (left) and unbounded (right).

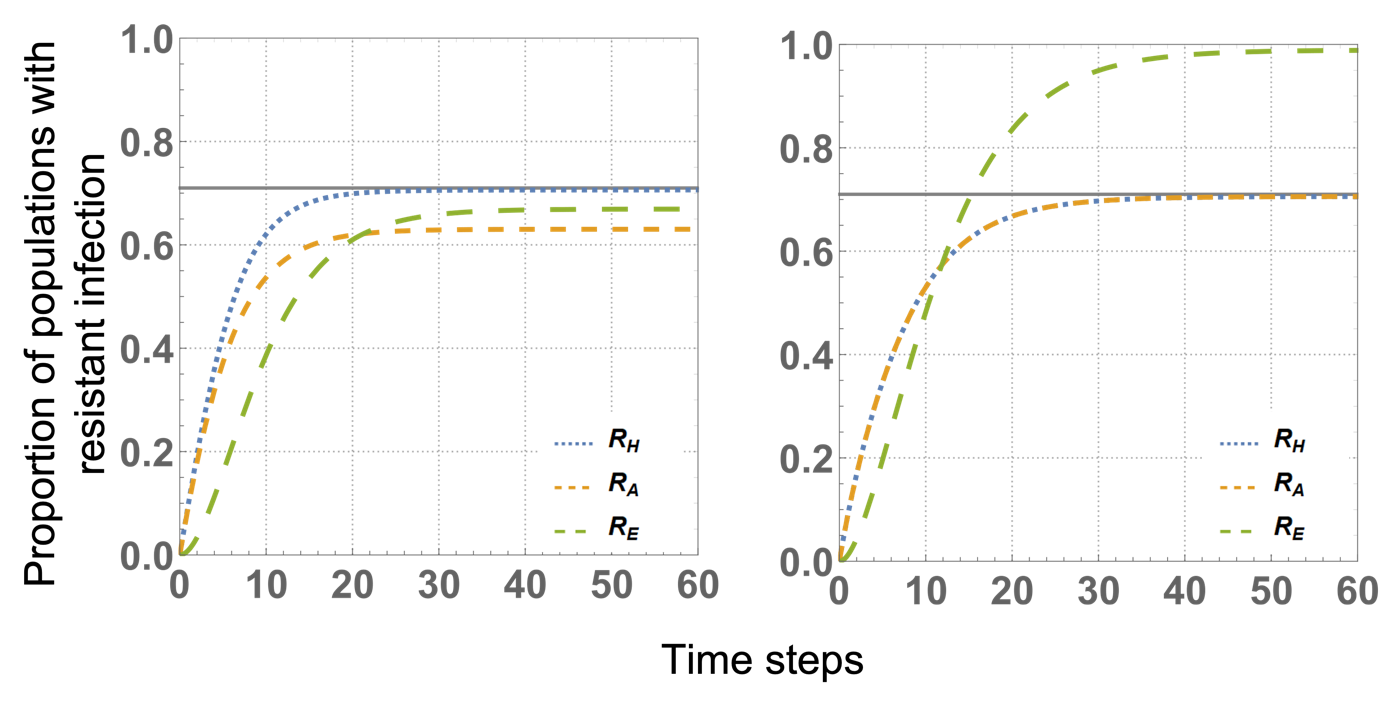

Figure 2: Heatmaps of the impact of reducing $\Lambda_{A}$, for different pre-intervention levels of $\Lambda_{A}$ (Y axis) and $\beta_{EH}$ (X axis), in all transmission scenarios.

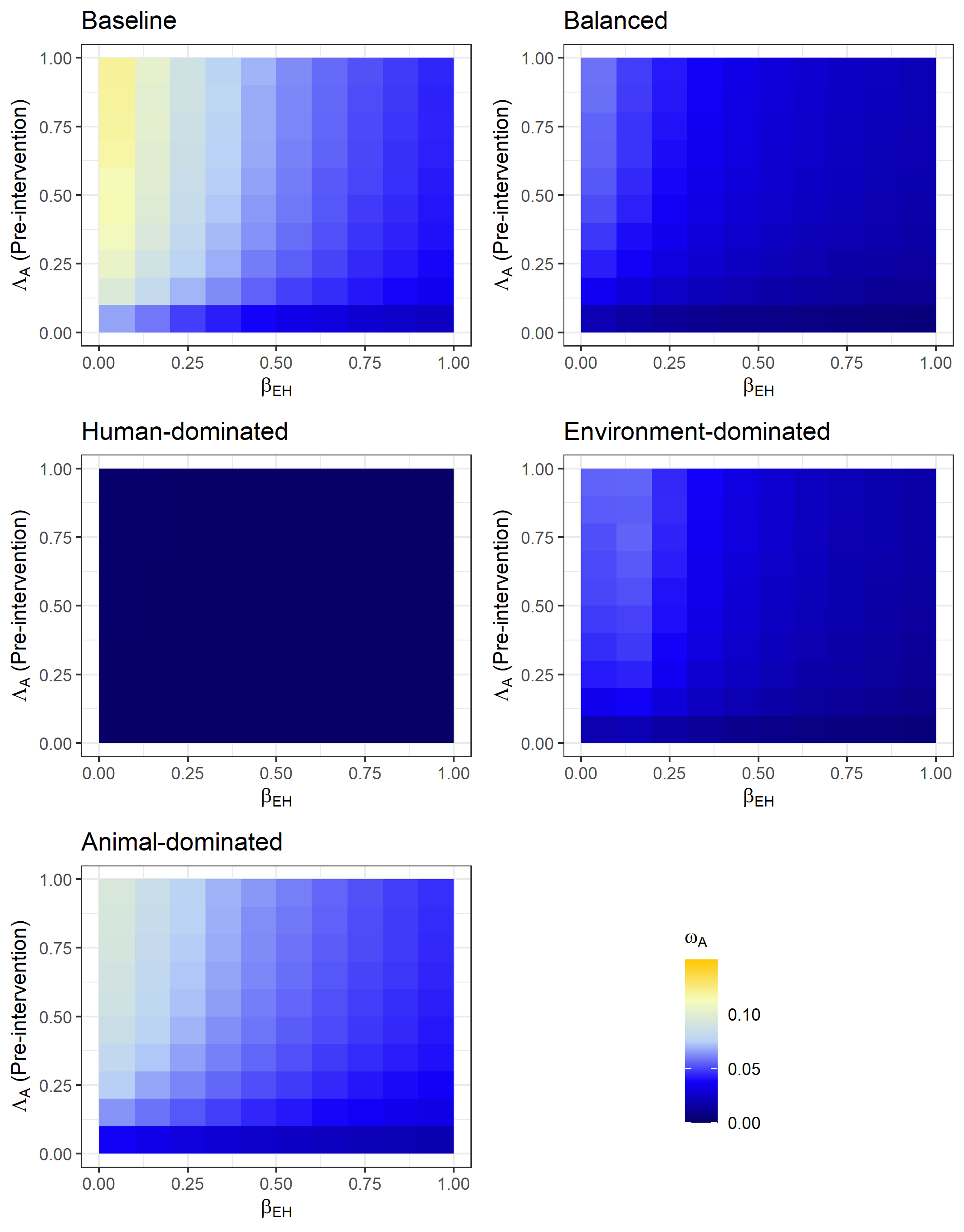

### Appendix references

EARS-Net. 2010. European Antimicrobial Resistance Surveillance Network.
